# Maturation of glutamatergic transmission onto dorsal raphe serotonergic neurons

**DOI:** 10.1101/2023.01.19.524776

**Authors:** Alexandre Kisner, Abigail M. Polter

## Abstract

Serotonergic neurons in the dorsal raphe nucleus (DRN) play important roles early in postnatal development in the maturation and modulation of higher order emotional, sensory, and cognitive circuitry. This unique position makes these cells a substrate by which early experience can be wired into brain. In this study, we have investigated the maturation of synapses onto dorsal raphe serotonergic neurons in typically developing male and female mice using whole-cell patch-clamp recordings in *ex vivo* brain slices. We show that while inhibition of these neurons is relatively stable across development, glutamatergic synapses greatly increase in strength between P6 and P21-23. In contrast to forebrain regions, where the components making up glutamatergic synapses are dynamic across early life, we find that the makeup of these synapses onto DRN serotonergic neurons is largely stable after P15. DRN excitatory synapses maintain a very high ratio of AMPA to NMDA receptors and a rectifying component of the AMPA response throughout the lifespan. Overall, these findings reveal that the development of serotonergic neurons is marked by a significant refinement of glutamatergic synapses during the first 3 postnatal weeks. This suggests this time as a sensitive period of heightened plasticity for integration of information from upstream brain areas and that genetic and environmental insults during this period could lead to alterations in serotonergic output, impacting both the development of forebrain circuits and lifelong neuromodulatory actions.

## Introduction

The formation and maturation of synapses is critical for proper development of fundamental behavioral processes (1). Disruption of synaptic development is linked to increased risk for neuropsychiatric illnesses such as anxiety disorders, autism spectrum disorders, substance use disorders and major depression (2–6). Understanding how synapses mature under standard developmental conditions allows us to define critical and sensitive periods in which environmental exposures and experience can lead to long-lasting alterations in the wiring and function of neuronal circuits. Developmental patterns of synapse assembly and maturation in projection neurons of forebrain regions such as the cortex and hippocampus have been well defined (7–9). It has become clear, however, that the course of synaptic maturation differs between cell types (1, 9), and considerably less is known about maturational processes in midbrain and hindbrain regions.

Serotonergic neurons within the brain’s dorsal raphe nucleus (DRN) play a vital role in brain development and influence a broad range of physiological processes as well as vulnerability to mental illnesses (2, 10). During early development, 5-HT acts as a trophic factor regulating cell division, differentiation, neuronal outgrowth, synaptogenesis and dendritic pruning (11–13). As development progresses, 5-HT neurons innervate cortical and subcortical areas, and the neurotrophic role of 5-HT is complemented by its neuromodulatory function. Precise levels of 5-HT signaling are required for structural and functional development of sensory (14, 15) and emotional circuits (16, 17), and genetic or pharmacological manipulation of 5-HT levels early in life results in structural and functional disruption of these circuits (18). Regulation of serotonergic neurons during early stages of life is of critical importance to proper experience dependent development of mature neuronal circuits. Despite this, little is known about how extrinsic inputs regulate 5-HT neurons over the course of maturation.

In rodents, 5-HT levels peak within the first postnatal week and then gradually decay until early adolescence and remain constant throughout adulthood (2). At postnatal day 4 (P4), DRN 5-HT neurons have a depolarized resting membrane potential and consequently increased excitability compared to later stages of development (19). Excitatory and inhibitory tone onto serotonergic neurons gradually increase across early postnatal development, although studies disagree about the precise timing of these events (19, 20). While 5-HT regulates circuit development in other brain areas, there is also ongoing maturation of synaptic inputs onto DRN 5-HT neurons. This suggests that early life experience could, through alterations in the developmental course of extrinsic inputs onto 5-HT neurons, orchestrate maturational changes across a wide range of neuronal circuits.

In this study, we use acute slice electrophysiology to investigate the maturation of synaptic properties of genetically-identified serotonergic neurons from the neonatal stage to adulthood. We show that the first three weeks are a time of dynamic change in the function and composition of excitatory synapses onto DRN 5-HT neurons. These synapses are predominated by AMPARs and at least a subset of these neurons express AMPARs with a high level of rectification, indicating persistence of calcium-permeable AMPARs (CP-AMPARs) into adulthood. These findings give new understanding of developmental processes in serotonergic neurons and provide an insight into windows of potential sensitivity to environmental perturbation.

## Materials and Methods

### Animals

All animals and experimental protocols were conducted in accordance with National Institutes of Health Guidelines for the Care and Use of Laboratory Animals, and with the approval of the IACUC of The George Washington University. Female and male ePet1-cre (Strain 12712, The Jackson Laboratory) crossed with Ai14 tdTomato reporter mice (Strain 7908, The Jackson Laboratory) (21, 22) transgenic mice were used in this study. Mice were group housed with littermates within ventilated cages in temperature- and humidity-controlled rooms with *ad libitum* access to water and rodent chow on a 12 h light/dark cycle.

### Electrophysiology

Mice were deeply anesthetized with ketamine (100 mg/kg) and dexmeditomidine (0.25 mg/kg) and perfused transcardially with ice-cold N-methyl-D-glucamine (NMDG)-based slicing solution (23) containing (in mM): 92 NMDG, 20 HEPES, 25 glucose, 30 NaHCO_3_, 1.2 NaH_2_PO_4_, 2.5 KCl, 5 sodium ascorbate, 3 sodium pyruvate, 2 thiourea, 10 MgSO_4_, and 0.5 CaCl_2_. Brains were rapidly dissected and placed in ice-cold NMDG solution. Horizontal brain slices (240 μm thick) containing the DRN were obtained using a vibratome (Leica VT1200, Leica Biosystems Inc., IL, USA). Immediately after slicing, brain slices were transferred to a holding chamber at 32°C degrees filled with a recovery solution containing (in mM): 92 NaCl, 20 HEPES, 25 glucose, 30 NaHCO_3_, 1.2 NaH_2_PO_4_, 2.5 KCl, 5 sodium ascorbate, 3 sodium pyruvate, 2 thiourea, 1 MgSO_4_, and 2 CaCl_2_. Slices were held at 32°C for one hour, and then at room temperature until use. For electrophysiological recordings, a single slice was transferred to a chamber perfused at a rate of 1.5 to 2.0 ml/min with heated (28-32°C) artificial cerebrospinal fluid (aCSF, in mM: 125 NaCl, 2.5 KCl, 1.25 NaH_2_PO_4_, 1 MgCl_2_ 6H_2_O, 11 glucose, 26 NaHCO_3_, 2.4 CaCl_2_. All solutions were saturated with 95% O_2_ and 5% CO_2_.

Fluorescent, tdTomato-positive neurons were located in brain slices. Whole-cell patch-clamp recordings were performed using a Sutter IPA amplifier (1 kHz low-pass Bessel filter and 10 kHz digitization) using Sutter Patch software (Sutter Instruments). Voltage-clamp recordings were made using glass patch pipettes with resistance 2-4 MOhms, filled with internal solution containing (in mM): 117 cesium methanesulfonate, 20 HEPES, 0.4 EGTA, 2.8 NaCl, 5 TEA-Cl, 2.5 Mg-ATP, and 0.25 Na-GTP, pH 7.3–7.4 and 285–290 mOsm. To measure the current-voltage relation of the AMPA-receptor component, spermine (0.1 mM) was added to the internal solution. Series resistance was monitored throughout voltage clamp recordings and recordings in which the series resistance changed more than 20% and/or exceed 20 MOhms were not included in the analysis. Membrane potentials were not corrected for junction potentials.

To measure the excitation-inhibition ratio, neurons were first voltage clamped at −70 mV, which approximates the reversal potential of GABAA receptors and allowed the exclusive detection of sEPSCs. Sequentially, the neurons were voltage clamped at 0 mV, the reversal potential for EPSCs, and sIPSCs were detected. Analysis of spontaneous PSCs was performed using SutterPatch software. A total of 220 synaptic events were detected from each cell using a threshold of 8 pA. The excitation-inhibition ratio for peak current was calculated by dividing the mean sEPSC peak current by the mean sIPSC peak current. Likewise, the excitation-inhibition ratio for frequency of synaptic events was determined by dividing the mean sEPSC frequency by the mean sIPSC frequency.

For analysis of evoked EPSCs, 100 μM picrotoxin was included in the aCSF. Electrical stimulation was carried out at 0.1 Hz using a bipolar stimulating electrode placed 100 – 200 μm rostral to the recording electrode. AMPAR/NMDAR ratio was calculated as the peak of AMPAR-mediated evoked-EPSC at −70 mV divided by the peak of the NMDAR-mediated evoked-EPSC at +40 mV after bath applying NBQX (10 μM) to isolate currents carried by NMDAR only. The respective evoked EPSC peak values were obtained by averaging at least 25 sweeps. The decay time constant (τ_decay_) of averaged AMPAR and NMDAR EPSCs were determined by fitting a single-exponential function using Igor-pro 9.0 based SutterPatch software (Sutter Instruments).

We determined the current-voltage relationship of AMPAR-mediated evoked EPSCs by measuring averaged (at least 25 sweeps) EPSC peak amplitude at 7 holding potentials between −70 mV and at +40 mV in the presence of the NMDAR antagonist AP-5 (50 μM) and spermine (100 μM) in the internal solution. The rectification index was calculated as the peak EPSC at +40 mV divided by the peak EPSC amplitude at −70 mV.

### Materials

All salts used for electrophysiology were purchased from Sigma-Aldrich (St. Louis, MO) or Fisher Scientific (Hampton, NH). Pharmacological reagents such as picrotoxin, spermine and tetrodotoxin were purchased from Tocris Biosciences (Bristol, United Kingdom). Ketamine and Dexmedetomidine were purchased from Covetrus (Elizabethtown, PA).

### Statistics

The results are reported as mean ± SEM. The number of cells and the number of animals used in each electrophysiological measurement are reported in the figure legends. Statistical tests were performed in GraphPad Prism 9.2 using one-way ANOVA and Tukey’s *post hoc* test was used for the comparisons within groups. Variance *F* values are reported in the text. In all analyses, results with a *P* < 0.05 were considered as significant.

## Results

### Maturation of spontaneous synaptic transmission

DRN 5-HT neurons were identified in brain slices by tdTomato fluorescence. To examine the maturation of spontaneous synaptic transmission onto DRN 5-HT neurons, we began by measuring spontaneous excitatory and inhibitory transmission over a developmental time course. Whole-cell recordings were made in acute brain slices from mice at the following time points across the juvenile, adolescent and adult period: P6, P15, P21-23, P33-37, P45-47 and P60-P80. We determined excitatory and inhibitory synaptic activity from the same DRN 5-HT neurons by voltage clamping cells at the reversal potential for GABAA (−70 mV) and ionotropic glutamate receptors (0 mV) and measured spontaneous excitatory postsynaptic currents (sEPSC) and spontaneous inhibitory postsynaptic currents (sIPSC), respectively.

We found that both sEPSCs and sIPSCs were readily apparent at P6 (**Fig. 1A**). The amplitude of sEPSC onto 5-HT neurons increased throughout the juvenile period, achieving a maximum at P21-23 and then remaining stable throughout adulthood (one way ANOVA, *F*_(5,70)_= 4.93, *p*=0.0006; **Fig. 1B**). The frequency of glutamatergic currents also showed an abrupt enhancement from P6 to P21-23, followed by a stabilization through adulthood (one way ANOVA, *F*_(5,67)_= 4.65, *p*=0.0011; **Fig. 1D**). Conversely, the amplitude of sIPSCs on 5-HT neurons did not significantly change across development (**Fig. 1C**), and the frequency of GABAergic postsynaptic currents only significantly increased from initial levels at P6 to P21-23 (one way ANOVA, *F*_(5,66)_= 2.66, *p*=0.029; **Fig. 1E**). These data suggest that glutamatergic transmission onto 5-HT undergoes significant maturation of presynaptic and postsynaptic function during the first three postnatal weeks, while inhibitory transmission remains relatively stable from the postnatal period through adulthood.

**Figure 1.**
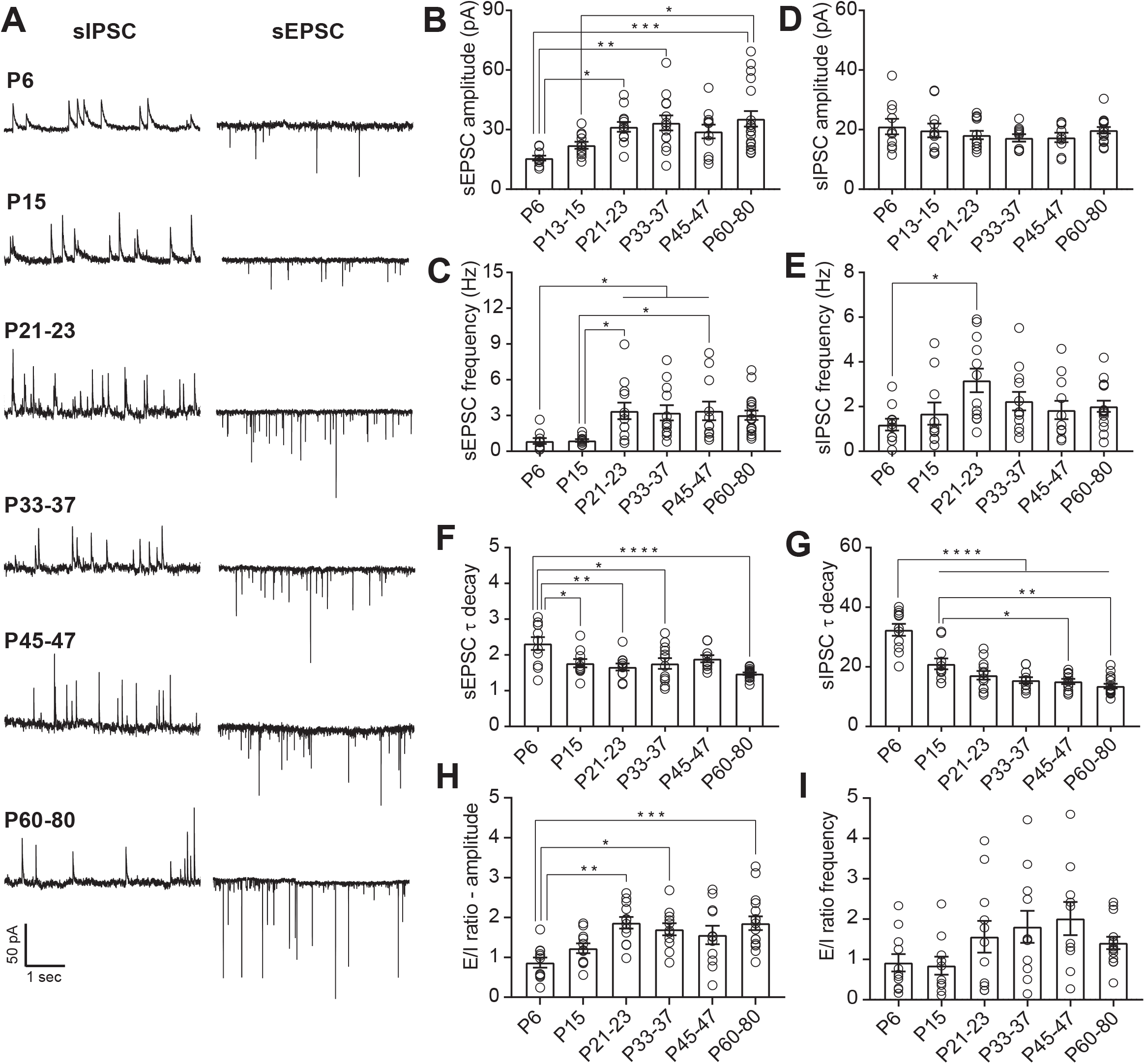
Maturation of spontaneous synaptic transmission onto DRN serotonergic neurons. **A:** Representative traces of spontaneous synaptic transmission at developmental timepoint. For each timepoint, traces were recorded from the same 5-HT neuron at holding potentials of −70 mV (sEPSC) and 0 mV (sIPSC). **B, C:** Developmental changes in the mean amplitude (B) and frequency (C) of sEPSCs. **D, E:** Developmental changes in the mean amplitude (D) and frequency (E) of sIPSCs. **F:** Time constant of decay (τ_decay_) for sEPSCs. **G:** τ_decay_ for sIPSCs. **H:** Excitation/Inhibition (E/I) ratio for amplitude of sPSCs. I: E/I ratio for frequency of sPSCs. PND 6, *n* = 10 cells (5 mice); PND 15, *n* = 11 cells (7 mice); PND 21-23, *n* = 12 cells (8 mice); PND 33-37, *n* = 12 cells (8 mice); PND 45-47, *n* = 11 cells (8 mice); PND 60-80, *n* = 17 cells (10 mice). Bar graphs represent mean ± SEM. \**p* < 0.05, one-way ANOVA followed by Tukey’s multiple comparison test.

As the subunit expression of postsynaptic glutamatergic and GABAergic receptors are known to change during development and consequently alter the kinetics of the postsynaptic currents, we next analyzed the decay time constant of the spontaneous postsynaptic currents. At P6, both sEPSC and sIPSC had slower decay time constant compared to later ages (one way ANOVA, *F*_(5,65)_= 6.37, *p* < 0.0001; **Fig. 1F**, one way ANOVA, *F*_(5,67)_= 26.95, *p* < 0.0001; **Fig. 1G**). The acceleration of sEPSC decay kinetics during maturation can primarily be attributed to factors such as a shift in subunit composition of AMPA receptors, the formation of more proximal somatic synapses and/or changes in synaptic structure (24–27). Similarly, the changes in sIPSC decay kinetics are most likely due to shifts in the composition of GABAA receptors (28). Maturation of synaptic tone across early postnatal life therefore occurs in concert with changes in the subunit composition of glutamatergic and GABAergic receptors.

The integration of convergent excitatory and inhibitory inputs plays a critical role in the maturation and stabilization of salient connections as well as in the regulation of neuronal output. Our data suggest that the maturation of the overall synaptic drive onto 5-HT is primarily determined by changes in glutamatergic transmission. To test this, we calculated the ratio of excitatory to inhibitory transmission onto 5-HT neurons across development. Analysis of the relationship between excitatory and inhibitory spontaneous neurotransmission indicate that at P6 and P15, the E/I ratio of current amplitude in 5-HT neurons is nearly equivalent. However, after this early postnatal period, excitation predominates and the E/I ratio peaks during adolescence (P21-23), remaining stable throughout adulthood (one way ANOVA, *F*_(5,67)_= 5.65, *p*=0.0002; **Fig. 1H**). In contrast to the E/I ratio of PSC amplitudes, no changes were seen in the ratio of E/I PSC frequencies. This indicates that developmental changes in E/I ratio are primarily driven by postsynaptic changes in glutamate receptor number or function.

### Excitatory currents onto 5-HT neurons are primarily mediated by AMPA receptors

Excitatory synaptic currents are primarily carried by AMPA and NMDA receptors, and the relative abundance of these receptors is developmentally regulated in many cell types (29). To examine developmental changes in the complement of synaptic glutamatergic receptors in 5-HT neurons we measured the AMPAR/NMDAR ratio. Across all developmental timepoints, serotonergic neurons exhibit a remarkably high AMPAR/NMDAR ratio, driven by the negligible presence of NMDA receptors. In contrast to what is typically seen in forebrain principal neurons (30–32), this AMPAR/NMDAR ratio was relatively stable across development, only showing a statistically significant difference between P6 and P45-47 neurons (one way ANOVA, *F*_(5,62)_= 2.64, *p*=0.03; **Fig. 2C**). These results indicate that across development, glutamatergic transmission onto 5-HT neurons is primarily mediated by AMPARs, with minimal contribution from NMDA receptors.

**Figure 2.**
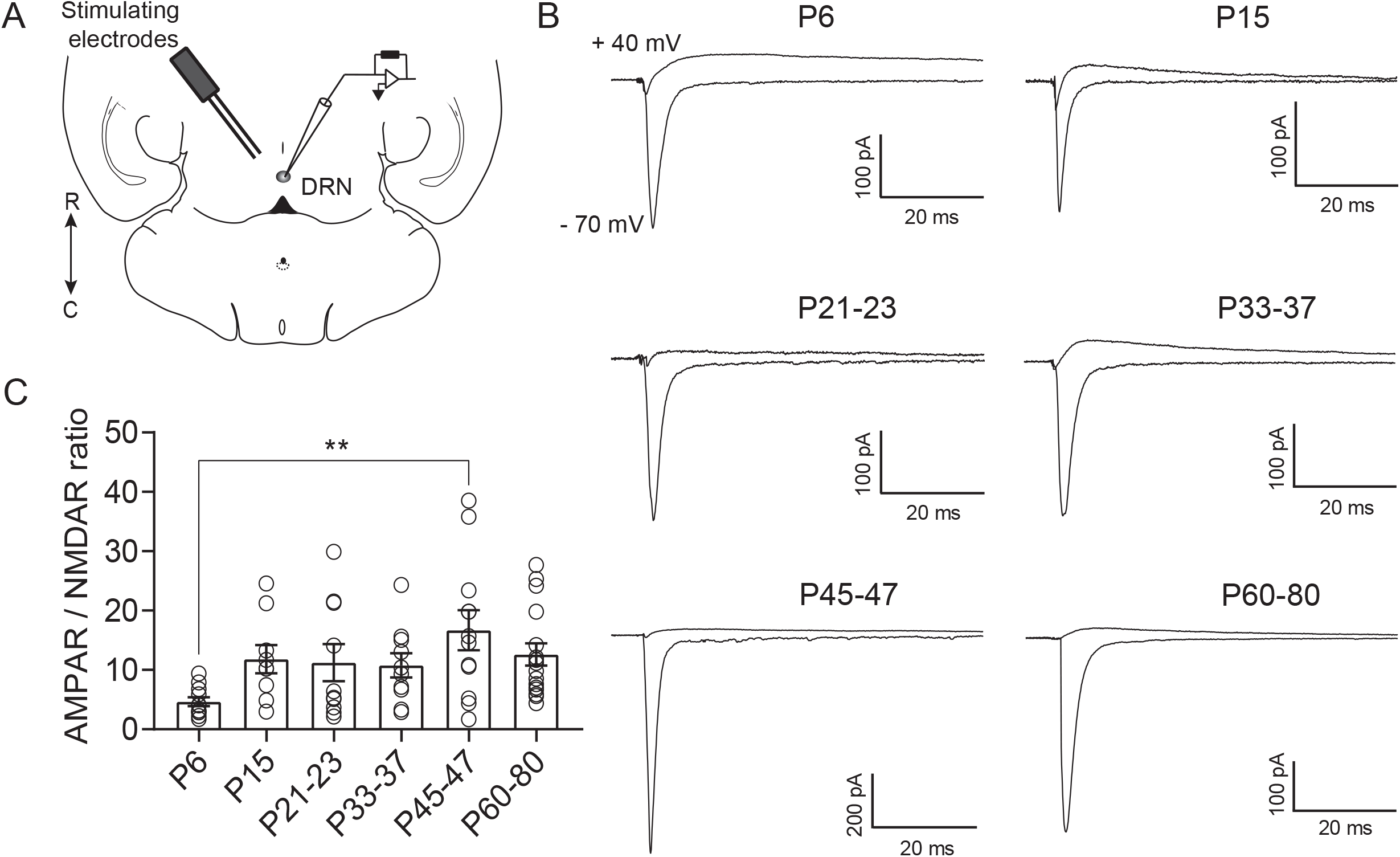
Excitatory currents onto DRN serotonergic neurons are primarily mediated by AMPAR. **A:** Schematic of the experiment illustrating the placement of the bipolar stimulating electrode in the DRN and recording of evoked excitatory currents from 5-HT neurons. **B:** Representative traces of electrically evoked excitatory synaptic currents recorded from 5-HT neurons at holding potentials of −70 mV and +40 mV. Recordings at +40 mV were performed in the presence of the AMPA antagonist NBQX (10 μM). **C:** Quantification of AMPA/NMDA ratios across maturation. PND 6, *n* = 11 cells (6 mice); PND 15, *n* = 9 cells (6 mice); PND 21-23, *n* = 10 cells (9 mice); PND 33-37, *n* = 10 cells (7 mice); PND 45-47, *n* = 12 cells (8 mice); PND 60-80, *n* = 16 cells (11 mice). Bar graphs represent mean ± SEM. ** *p* < 0.01, one-way ANOVA followed by Tukey’s multiple comparison test.

Both AMPARs and NMDARs have multiple subunits which are dynamically regulated over development in forebrain regions. Subunit composition alters the decay kinetics of both AMPARs and NMDARs (33–36). To investigate this, we examined the τ_decay_ of AMPAR and NMDAR-mediated evoked EPSCs. We found that the τ_decay_ constant of AMPAR-mediated EPSCs across maturation decreased between P6 and P33-37 (one way ANOVA, *F*_(5,59)_=5.96, *p* < 0.0001, **Figs. 3A-B**). The decay time of the NMDAR-mediated EPSC was also significantly slower in P6 neurons than at later ages (one way ANOVA, *F*_(5,53)_= 9.47, *p* < 0.0001; **Figs. 3C-D**). Similar to our findings from spontaneous EPSCs, these results suggest that both types of receptors undergo changes in subunit composition early in development that may contribute to the stabilization of glutamatergic synapses.

**Figure 3.**
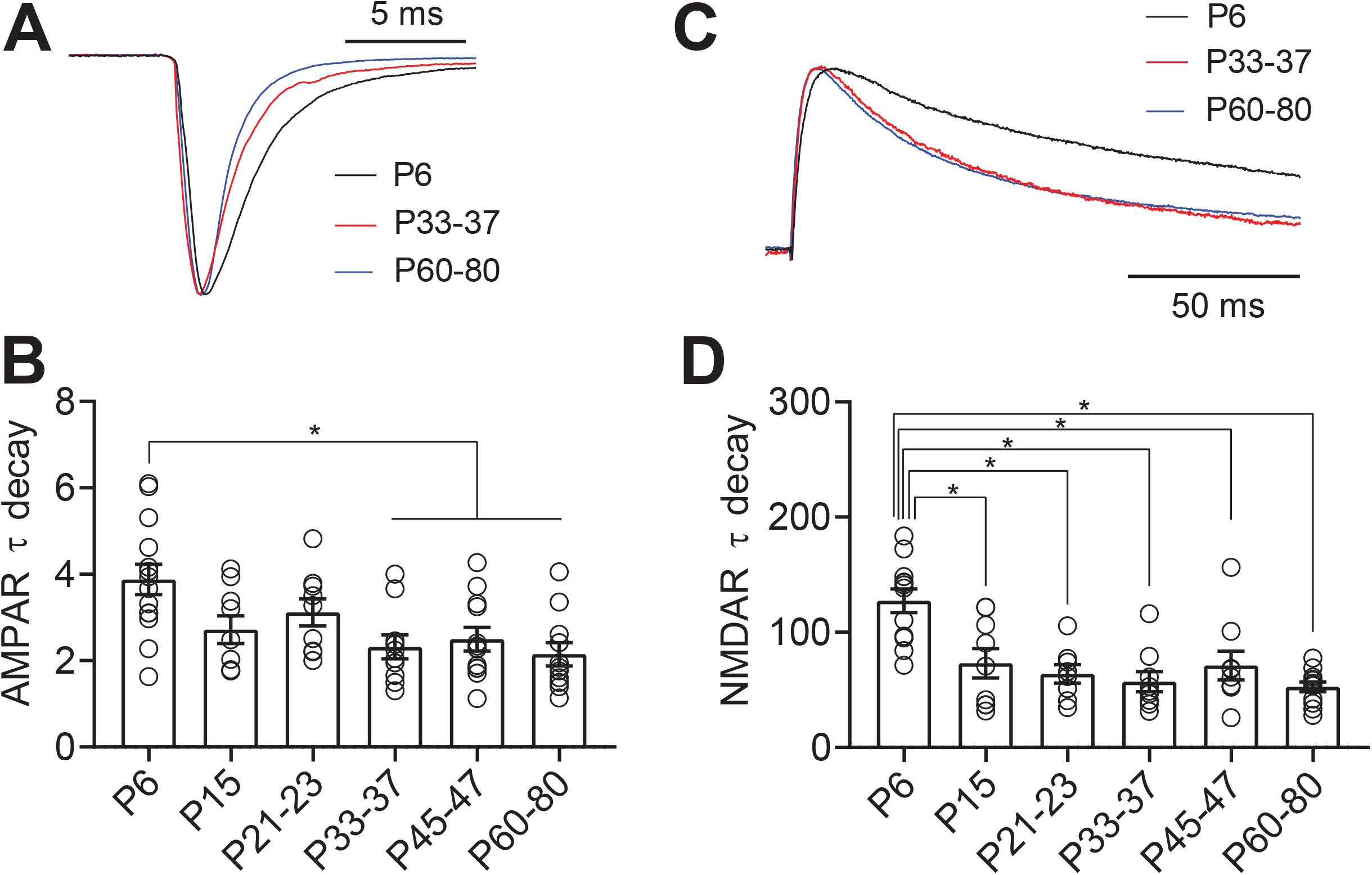
AMPA and NMDA receptor subunits are developmentally regulated in 5-HT neurons. **A:** Scaled representative traces of evoked AMPAR currents. **B:** Time constant of decay (τ_decay_) for evoked AMPAR currents recorded at −70 mV. **C:** Scaled representative traces of evoked NMDAR currents. **D:** Time constant of decay (τ_decay_) for evoked NMDAR currents recorded at +40 mV in the presence of the AMPA antagonist NBQX (10 μM). Bar diagrams represent mean ± SEM. * *p* < 0.05, one-way ANOVA followed by Tukey’s multiple comparison test.

### 5-HT neurons express AMPARs that are inwardly rectifying into adulthood

At many forebrain excitatory synapses, GluA2-lacking, CP-AMPARs predominate in early development and are gradually replaced by GluA2-containing receptors by the onset of adolescence. These shifts in receptor composition are essential to the establishment of critical periods and age-dependent mechanisms of experience-dependent plasticity. In these cells, reinsertion of CP-AMPARs in adult animals can also be an important mechanism of plasticity underlying learning and substance abuse (37–39). However, this pattern of maturation is not universal, as excitatory synapses onto other cell types, such as parvalbumin positive interneurons in the cortex and hippocampus, maintain CP-AMPARs as the predominant receptor subtype throughout the lifespan (40–43). We next examined whether serotonergic neurons likewise exhibit shifts in the prevalence of CP-AMPARs across the lifespan. As CP-AMPARs exhibit strong inward rectification (44–46), rectification of AMPAR-mediated currents is used as an indicator of the presence of CP-AMPARs at a synapse. Therefore, we pharmacologically isolated and examined the voltage dependence of evoked AMPAR-mediated synaptic currents (**Fig. 4A**). We found that AMPAR-mediated currents onto 5-HT neurons exhibited a largely stable rectification index across development and did not show replacement of CP-AMPARs with calcium-impermeable AMPARs in the early postnatal period (**Fig. 4B**). The only significant change in rectification was apparent in P21-23 neurons, which exhibited a significantly higher level of rectification than other timepoints. This suggests that at this time point glutamatergic synapses onto 5-HT neurons express either a much lower or negligible proportion of CP-AMPA receptors (**Fig. 4C**). Intriguingly, this shift in rectification is highly specific to this timepoint, indicating a transient change in the composition of excitatory synapses. We also found that the rectification index was highly heterogeneous within each timepoint-some cells exhibiting almost complete rectification, while others showed almost none. Thus, while most cells express some level of CP-AMPARs throughout development, the proportion of receptors made up of these subunits varies from cell to cell. This may reflect heterogeneity of subtypes of serotonergic cells in the DRN, or slice-to-slice variation in the inputs sampled by electrical stimulation.

**Figure 4.**
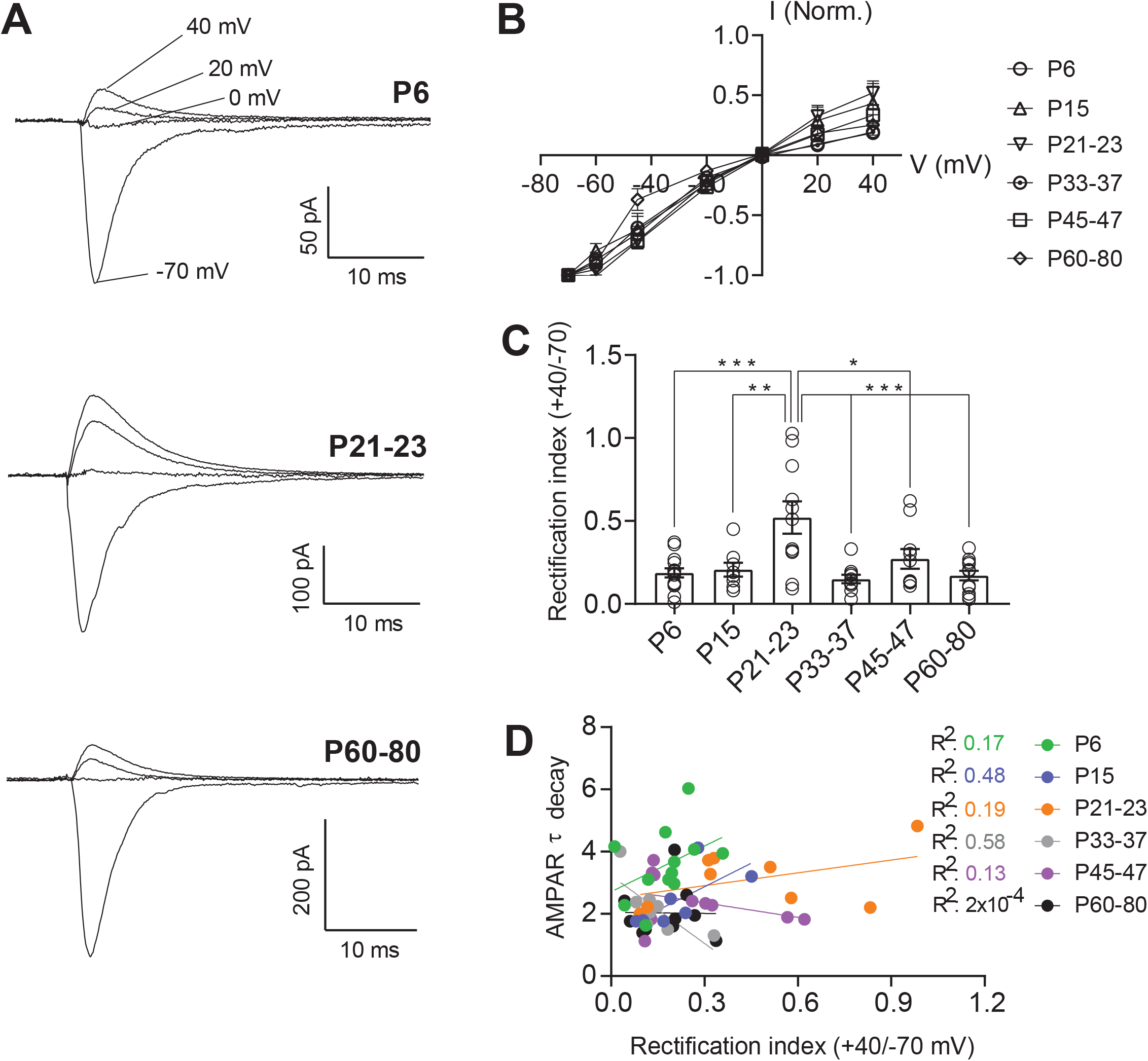
AMPAR-mediated currents are inwardly rectifying throughout development in a subset of 5-HT neurons **A:** Representative traces of evoked excitatory synaptic currents recorded from 5-HT neurons at holding potentials of −70 mV to +40 mV. **B:** Average peak current – voltage curves with slight deviation from a linear relationship. **C:** Rectification index of AMPAR currents, calculated as the ratio of peak amplitude at +40 mV divided by the peak amplitude at −70 mV for each cell. **D:** Correlation between AMPAR EPSC τ decay and rectification index for each time point. PND 6, *n* = 14 cells (7 mice); PND 15, *n* = 8 cells (7 mice); PND 21-23, *n* = 11 cells (6 mice); PND 33-37, *n* = 10 cells (5 mice); PND 45-47, *n* = 10 cells (7 mice); PND 60-80, *n* = 12 cells (7 mice). Bar graphs represent mean ± SEM. \**p* < 0.05, one-way ANOVA followed by Tukey’s multiple comparison test.

**Figure 5.**
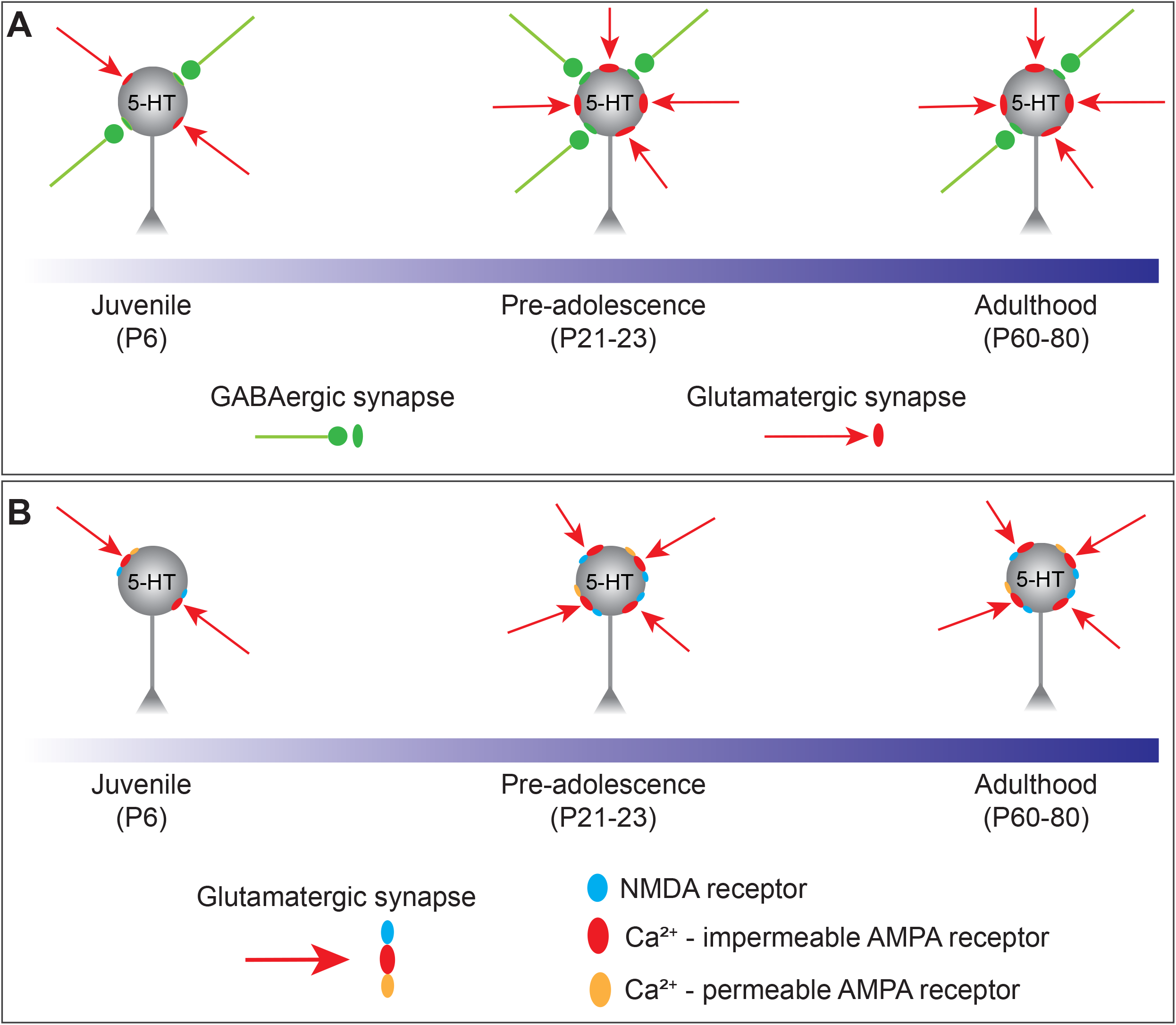
Schematic summarizing the maturation of synaptic inputs onto 5-HT neurons. **A:** The overall synaptic drive is led by glutamatergic transmission, which undergoes an abrupt enhancement between juvenile and the pre-adolescent period and then remained stable throughout adulthood. **B:** Glutamatergic transmission is mainly carried by AMPA receptors and in a subset of 5-HT neurons these receptors have a high permeability to calcium and are inwardly rectifying across the whole developmental period.

As shown in Figure 3A-B, AMPAR-mediated currents show increases in the τ_decay_ between P6 and P33-37. While CP-AMPARs exhibit faster kinetics than calcium-impermeable AMPARs, we hypothesized that the relative stability of the rectification index over this time period indicated that changes in τ_decay_ were likely independent of CP-AMPARs. To test this, we performed an analysis of the correlation between τ_decay_ and rectification index of AMPAR-mediated currents within the same cells (**Fig. 4D**). We found no significant correlation between these metrics, indicating that as expected, the developmental changes in AMPAR τ_decay_ kinetics are not related to fluctuations in the prevalence of CP-AMPARs.

## Discussion

### Developmental dynamics of DRN synaptic function

DRN 5-HT neurons undergo significant maturation during the initial postnatal period. These cells exhibit high levels of excitability shortly after birth, which rapidly declines until reaching stable levels at P21-23 (19). As synaptic inputs are a significant driver of cellular excitability, and a key site for the contribution of environmental and sensory experiences to the development of circuits, we sought to characterize the maturation of excitatory and inhibitory synapses onto 5-HT neurons. Prior studies addressing this question have yielded somewhat conflicting information. In one study, using ePet-YFP reporter mice, both sEPSC and sIPSC amplitudes were stable across development from P4-P60, while sEPSC and sIPSC frequency gradually increased (19). In contrast, in a study recording from VGAT-negative (putative serotonergic) neurons in the DRN, investigators found that both amplitude and frequency of sEPSCs and sIPSCs decreased between P5-7 and P15-17 (20).

Given this uncertainty, we therefore first investigated the maturation of excitatory and inhibitory synaptic tone onto serotonergic neurons in ePet-Cre-tdTomato mice. We find that excitatory transmission increases in strength between P6 and P21-23. Intriguingly, post-synaptic strength, as measured by sEPSC amplitude, increases gradually across this time period, while presynaptic strength increases abruptly between P15 and P21-23. Prior work has shown that this third postnatal week coincides with an expansion of dendritic branching of serotonergic neurons (19); it is therefore likely that this increase in frequency represents an increase in the number of excitatory synaptic inputs onto serotonergic neurons. Although the maturation of synapses is governed by different factors, the postsynaptic expression and activation of specific serotonergic receptors such as 5-HT4 on 5-HT neurons is likely to play a pivotal role in the formation and maturation of glutamatergic synapses (47). Taken together, this suggests that during the first three weeks of postnatal life, changes in postsynaptic responsivity and number of synaptic sites occur at distinct timescales to contribute to growing strength of excitatory inputs onto DRN serotonergic neurons.

In contrast to the dynamic changes in excitatory transmission, sIPSC amplitude and frequency are largely stable between birth and adulthood. This suggests that the number and strength of inhibitory synapses is set shortly after birth and that changes in responsivity to incoming stimuli across postsynaptic development are mediated by addition and functional changes of excitatory inputs. We calculated the E/I ratio for both PSC frequency and amplitude to examine the balance of excitation and inhibition within individual cells across maturation. Intriguingly, we find an increase in the E/I ratio of PSC amplitude, but not frequency, between P6 and P21-23, followed by stability across adolescence and adulthood. This supports a model by which inhibitory tone is established early in postnatal life and exerts a constant influence on DRN 5-HT neurons, while excitatory synapses increase in number and strength as the animal encounters increasingly complex sensory, social, and emotional stimuli throughout the first three weeks of postnatal life.

Across several studies, there is broad agreement that the majority of synaptic maturation onto DRN serotonergic neurons occurs in the first three weeks of life. However, the specifics of this maturation vary across labs. There are several potential explanations for differing patterns of synaptic maturation across multiple studies. Our work and the two prior studies used three different mouse lines to identify serotonergic neurons, introducing potential variability in genetic background as well as in neurons labelled by specific genetic strategies (19, 20). There are also differences in slice preparation, as our study was performed in horizontal sections and prior studies were conducted in coronal sections. This likely results in differences in the preservation of inputs from distal and local sources, and in which inputs are sampled by electrical stimulation. This highlights a potential heterogeneity in developmental processes across distinct inputs, suggesting there is much exciting data to be found in future studies of input-specific synaptic maturation. Finally, developmental processes are experience dependent. Differences in housing conditions and husbandry practices between institutions may lead to subtle differences in developmental patterns between studies.

The dynamic changes in synaptic strength highlight the first three weeks as an important developmental period for DRN serotonergic neurons. This timing coincides with maximal developmental synaptogenesis in other brain areas of rodents, including the primary visual cortex, barrel, motor, frontal and somatosensory cortex (48–50). The timing of maturation and stabilization of glutamatergic synapses in the DRN also coincides with the anatomical maturation of serotonergic innervation of the forebrain, as serotonergic axons generally reach their postsynaptic targets by the end of the third postnatal week (51). This close temporal relationship between the maturation of afferent inputs and the serotonergic arborization in other brain areas suggests a synchrony in these processes. Given the role that serotonin plays in maturation and stabilization of sensory and affective circuits (14–17), genetic or environmental insults that disrupt maturation of synapses onto DRN serotonergic neurons may have wide-ranging developmental ripples throughout a number of behavioral domains.

### Dorsal raphe glutamatergic receptors across development

Fast glutamatergic synaptic transmission is primarily mediated by AMPA and NMDA receptors, and maturation of forebrain synapses is associated with changes in the relative ratio of these two receptors, as well as the composition of subunits making up both receptor types (52, 53). In forebrain excitatory neurons, more mature synapses typically have a higher ratio of AMPA to NMDA receptors. In addition, the subunit composition of AMPA receptors shifts across development, as GluA2-lacking, CP-AMPAR are replaced by GluA2-containing non-CP-AMPARs. However, as genetic tools have increasingly allowed the investigation of synaptic development onto specific cell types, it has become clear that the process of synaptic development and composition of mature synapses varies significantly by cell type. For example, in the hippocampus and cortex, some subtypes of interneurons exhibit minimal NMDAR-mediated excitatory transmission and the major glutamatergic synaptic response is mediated by CP-AMPARs throughout life (40, 42, 54, 55). We find that DRN 5-HT neurons exhibit remarkable stability of AMPA/NMDA ratios across development. These neurons exhibit high ratios of AMPA/NMDA receptors compared to many cell types, an effect that is driven by small evoked NMDAR-mediated currents. This suggests that DRN 5-HT neurons express a very low level of NMDA receptors at excitatory synapses throughout life. Given the critical role that NMDA receptors play in experience-dependent synaptic plasticity (56, 57), future work investigating mechanisms of plasticity at these synapses will be of high interest.

In addition to exhibiting a high, stable ratio of AMPA receptors to NMDA receptors across development, AMPAR-mediated currents onto DRN 5-HT neurons exhibit a robust level of rectification across the lifespan. This indicates that these receptors likely contain low levels of GluR2 subunits and have high calcium permeability (36). During early development in forebrain areas, CP-AMPAR are abundant in many synapses, but as the neurons mature levels of these receptors diminish and calcium-impermeable AMPAR become dominant (33, 58, 59). However, in distinct neuronal populations of interneurons in a number of brain regions, CP-AMPARs are predominant even in adulthood (40–43). AMPA receptors on DRN 5-HT neurons exhibit a largely unchanged level of rectification across development, suggesting that some proportion of glutamatergic synapses onto these cells express calcium permeable AMPA receptors throughout life. This is consistent with prior work in rats showing a residual sensitivity of AMPAR-mediated currents to the CP-AMPAR antagonist NASPM, even in adults (60). While the functional implications of the presence of calcium permeable-AMPARs in 5-HT neurons are unclear, calcium-impermeable and CP-AMPARs have distinct roles in integrating excitatory transmission for maturation of synaptic strength and plasticity (61). Given the low levels of NMDA receptors in DRN 5-HT neurons, it is intriguing to speculate that calcium influx through CP-AMPARs may play a critical role in producing and sustaining plastic changes (41, 43).

Intriguingly, AMPARs onto 5-HT neurons undergo a brief, transient increase in the expression of calcium impermeable AMPARs at P21-23. This timeframe corresponds to a period in which dendritic branching of 5-HT neurons expands, and which excitatory tone onto 5-HT neurons reaches its peak (19). This suggests a potential model by which newly added synapses in this time period express calcium impermeable AMPARs, which are replaced by synapses or receptors containing CP-AMPARs as adolescence progresses. Thus, although excitatory tone remains stable, more subtle changes marked by shifts in AMPAR composition refine function at excitatory synapses during the early phase of adolescence.

Our data suggests that there are additional maturational changes in the complement of receptors available at synapses. The decay kinetics of AMPAR-mediated EPSCs become faster during maturation, consistent with prior work in auditory and cerebellar granule cell synapses (24, 25, 27, 62). Several factors influence the decay time of AMPAR-mediated EPSCs, including changes in the AMPAR subunit composition of the receptor, dynamics of glutamate transporters and physical structure of the synapse. Our analysis indicates that the changes in decay kinetics do not correlate with rectification index within a given cell, telling us that presence of CP-AMPARs is not what determines decay constant in these cells. However, it is possible that other subunit changes may underlie this alteration in receptor kinetics. For instance, in the calyx of Held, an AMPAR subunit change from slow-gating GluR1 to fast-gating GluR3/4 plays a functional role in maintaining high-fidelity fast neurotransmission (24).

The decrease in NMDAR-mediated synaptic current decay kinetics between P6 and later ages also suggests a developmental switch in NMDAR subunit composition, likely from NR2B to NR2A-containing receptor as observed at other synapses (63–67). Given the critical role of NMDAR containing NR2B subunits in immature glutamatergic synapses controlling AMPAR expression, developmental changes in NMDAR subunit composition may also regulate the maturation of excitatory transmission and synaptic plasticity in 5-HT neurons (68). For example, the transiently slower decay time kinetics of NMDAR containing NR2B during early postnatal period may provide higher calcium influx that would contribute for a window of heightened plasticity for 5-HT neurons (30). However, as neurons mature, NR2B subunits are replaced by NR2A containing NMDARs (68). The faster decay time kinetics of NR2A containing NMDARs prevents overload of calcium influx and helps to stabilize the maturation of the synapse.

These results suggest that dorsal raphe excitatory synapses undergo changes in synaptic strength and receptor makeup over development. However, it is important to remember that these represent global measurements across all ePet-Cre+ serotonergic neurons and synaptic inputs. Emerging evidence tells us that serotonergic neurons are not a monolithic block, but rather are made up of several unique subtypes with differing behavioral roles, projection targets, and neurotransmitter content (69). Likewise, excitatory and inhibitory inputs to DRN serotonergic neurons arise from many sources, and different synaptic inputs may exhibit different properties and developmental patterns (70–72). While modern viral optogenetic and anatomical tracing tools have allowed a revolution in study of input and output specific plasticity, it remains extremely challenging to apply these tools in deep brain regions of juvenile animals. Rising to this challenge, as well as development of less invasive tools to label and manipulate increasingly specific cell groups, will be critical to truly understand the developmental landscape of the DRN.

### Functional implications

Activity of DRN serotonergic neurons is driven by an integration of both excitatory and inhibitory synaptic inputs from local and distal neurons (70–72), and development of the serotonergic system is regulated by coordinated changes in synaptic connectivity and function. Our findings here describe the maturation of glutamatergic synaptic transmission onto 5-HT neurons extending previous work demonstrating that morphological, anatomical, and physiological functions of these cells are dynamic throughout development (19, 20, 73). Functional changes in the ability of external stimuli to drive activation of 5-HT neurons is particularly critical across development given serotonin’s dual neurotrophic and neuromodulatory roles. Changes in responsivity of serotonergic neurons will not only shape ongoing behavior of developing animals but will also sculpt the structure and function of the forebrain circuits that receive serotonergic innervation.

The three first postnatal weeks represent an important period for formation and stabilization of excitatory inputs onto serotonergic neurons, and thus a period in which they are likely to be particularly sensitive to perturbation. That this phase of synaptic development appears to stabilize at P21-23 is intriguing, as in our lab and many others weaning occurs at this timepoint. This suggests that serotonergic neurons are highly excitable specifically during an early time period where juveniles are protected by and learning from parental influence, and transition to a less excitable, more stable state upon leaving life in the nest. Early life experiences such as quality or duration of parental care, enrichment or poverty of environmental influences, or exposure to numerous chemical or dietary factors are likely to shift this maturational process, potentially leading to downstream physiological and behavioral effects. A fuller knowledge of how this system matures could therefore have significant implications for our understanding of developmentally-rooted affective disorders such as anxiety, depression, and autism spectrum disorders. Future studies investigating how early life experiences shape maturation of the dorsal raphe and serotonergic circuits will be particularly valuable.

## Acknowledgements

The authors thank Ms. Caitlyn Cody and Ms. Brittney Thompson for assistance with mouse husbandry and genotyping. This work was supported by NIMH R00106757, a research grant from the Margaret Q. Landenberger Foundation, and award number UL1TR001876 from the NIH National Center for Advancing Translational Sciences.

